# Long-term cross-scale comparison of grazing and mowing on plant diversity and community composition in a salt-marsh system

**DOI:** 10.1101/2021.03.12.435073

**Authors:** Qingqing Chen, Jan P. Bakker, Juan Alberti, Elisabeth S. Bakker, Christian Smit, Han Olff

## Abstract

1. Land abandonment is increasing in recent decades in Europe, usually accompanied by a decline in biodiversity. Whether livestock grazing and mowing can safeguard biodiversity across spatial scales in the long term is unclear.
2. Using a 48-year experiment in a salt marsh, we compared land abandonment (without grazing and mowing) and seven management regimes including cattle grazing, early season mowing, late season mowing, both early and late season mowing, and grazing plus each of the mowing regimes on plant diversity at the local (i.e. plot) and landscape scales (i.e. across plots). Also, we compared their effects on community composition (both in identities and abundances) in time and space.
3. Under land abandonment, plant diversity declined in the local communities and this decline became more apparent at the landscape scale, particularly for graminoids and halophytes. All management regimes, except the late season mowing, maintained plant diversity at these scales.
4. Local plant communities under all treatments underwent different successional trajectories, in the end, diverged from their initial state except for that under grazing (a cyclic succession). Interannual composition change remained stable over time under land abandonment and grazing plus early season mowing. It increased over time under grazing and late season mowing, it increased in the second half of the experiment under other treatments. Vegetation homogenized in the landscape over time under land abandonment while vegetation was heterogeneous under all management regimes.
5. *Synthesis.* Our experiment suggests that late season mowing may not be sustainable to conserve plant diversity in salt marshes. Other management regimes can maintain plant diversity across scales and vegetation heterogeneity in the landscape in the long term, but local community composition may change over time.

## Introduction

Food production has been increasingly intensified in high-productive farming systems, while low-productive farming systems that harbor high biodiversity have been abandoned (Bignal & McCracken, 1996; Terres et al., 2015; Ustaoglu & Collier, 2018). In Europe, land abandonment has been widely observed in grasslands and salt marshes(Bakker, Bos, & De Vries, 2003; Poschlod, Bakker, & Kahmen, 2005). Grasslands and salt marshes have striking differences (salt marshes face salinity and inundation stress; Bakker, 1989), but they also share similarities. For instance, both systems are dominated by graminoids and traditionally grazed by livestock for food production (e.g. meat and milk). Similarly, mowing for hay (to feed livestock) used to be a common practice in grasslands and salt marshes (Bakker, 1989; Gedan, Silliman, & Bertness, 2009; Poschlod et al., 2005). Mowing is still common in grasslands, but it is rare in salt marshes (Esselink, 2017; Tälle et al., 2016). Land abandonment in grasslands and salt marshes is usually accompanied by negative consequences such as a decline in biodiversity (Terres et al., 2015; Ustaoglu & Collier, 2018).

To reverse the trend of biodiversity decline, herbivore reintroduction gained popularity in recent years (Garrido et al., 2019; Kumm, 2003). However, whether livestock grazing is optimal to conserve plant diversity in the long term is still debated (Bakker et al., 2003). Although livestock grazing generally increases plant diversity in European grasslands and salt marshes (Davidson et al., 2017; Tälle et al., 2016), these results are usually based on shorter-term experiments (< 15 years). In the long term (time scale across multiple decades), herbivores may promote dominance of gazing-tolerant or resistant plant species (Olff & Ritchie, 1998), which in turn may decrease plant diversity (Koerner et al., 2018). Alternatively, herbivores may promote habitat heterogeneity over time by trampling and localized deposition of urine and dung (Gillet, Kohler, Vandenberghe, & Buttler, 2010; Ludvíková, Pavlů, Gaisler, Hejcman, & Pavlů, 2014; R. Van Klink et al., 2015), a heterogeneous environment usually promotes plant diversity (Davies et al., 2005; Lundholm & Larson, 2003). Meanwhile, whether other traditional management tools such as mowing can be useful alternatives to livestock grazing in conserving plant diversity in the long term remains elusive. To our knowledge, the effects of livestock grazing and mowing on plant diversity in salt marshes have been rarely compared (but see Bakker, 1978). Shorter-term experiments comparing grazing and mowing in grasslands yielded inconsistent results (e.g. Catorci, Cesaretti, Malatesta, & Tardella, 2014; De Cauwer & Reheul, 2009; Wellstein, Otte, & Waldhardt, 2007), while longer-term experiments are rare. Therefore, comparisons of long-term livestock grazing and mowing are needed to gain insight into improved management in conserving diversity in abandoned land (Garrido et al., 2019; Tälle et al., 2016).

Effects of grazing and mowing on plant diversity and community composition may be different (Tälle et al., 2016), and these differences may increase over time. Herbivores remove aboveground biomass continuously and selectively (Tälle *et al.* 2016). Mowing also removes aboveground biomass, although it removes biomass drastically and uniformly. Moreover, herbivore trampling and deposition of urine and dung can also impact plant diversity and community composition (Kobayashi, Hori, & Nomoto, 1997; Kohler, Gillet, Gobat, & Buttler, 2004; Lezama & Paruelo, 2016; Ludvíková et al., 2014). Therefore, the effects of herbivore grazing may differ from that of mowing on plant diversity, particularly on functional groups. For instance, herbivores may suppress forbs while promoting graminoids because graminoids have basal meristems which may be more resistant to trampling (Briske & Richards 1995). Also, tiller-forming graminoids can tolerate repeated biomass removal better via rapid compensatory growth (Van Der Graaf, Stahl, & Bakker, 2005). Trampling may induce anoxic conditions in soil similar to that of inundation, which may promote halophytes (Van Klink *et al.* 2015). Halophytes are plant species well adapted to saline and anoxic conditions (Scherfose 1990; Bakker, Esselink, Dijkema, Van Duin, & De Jong, 2002). These trampling effects tend to accumulate over time (Elschot, Bakker, Temmerman, Van De Koppel, & Bouma, 2015; Mikola et al., 2009; R. Van Klink et al., 2015). Therefore, the effects of herbivore grazing and mowing on plant diversity and community composition may become more distinct in the long term (Chen, Bakker, Alberti, & Smit, 2020; Moog et al., 2002).

Not only long-term but also cross-scale monitoring is needed to fully evaluate biodiversity changes in response to ecological drivers such as grazing and mowing (Chase et al., 2018) and to inform conservation and management (Isbell et al., 2017). However, ecologists often focus on small spatial scales (plots) (Collins, Knapp, Briggs, Blair, & Steinauer, 1998; De Cauwer & Reheul, 2009; Kahmen, Poschlod, & Schreiber, 2002; Lepš, 2014). Studies show that the effects of herbivore grazing and mowing on plant communities are sometimes scale-dependent (Collins, Glenn, & Briggs, 2002; Lepš, 2014; Oldén & Halme, 2016; Wanner et al., 2014). For instance, Wanner et al., (2014) found that the effects of herbivores on plant diversity are more apparent in the local communities (4 m^2^) than at the landscape scale in a Wadden Sea salt marsh. However, Oldén and Halme (2016) found that herbivores not only increase plant diversity in the local communities but also promote vegetation heterogeneity in the landscape in a woody pasture in Finland. A heterogeneous vegetation structure at a landscape scale is often needed to warrant biodiversity at multiple trophic levels (Roel van Klink et al., 2016; Wanner et al., 2014). To our knowledge, the long-term effects of grazing and mowing on plant diversity and community composition at multiple spatial scales have rarely been compared.

Here, using a 48-year experiment in a salt marsh, we compare the effects of land abandonment (control; without grazing and mowing), and seven management regimes on plant diversity and community composition across spatial scales. These management regimes include cattle grazing, early season mowing, late season mowing, both early and late season mowing, and grazing plus each of the mowing regimes. Specifically, we ask two questions: (1) how do these eight treatments impact plant diversity and diversity in different functional groups at the local (i.e. plot) and landscape scales (across plots); (2) how do treatments alter community composition in time and space (local and landscape scale, respectively).

## Materials and Methods

### Study system

The experiment was conducted in the natural salt marsh of the barrier island of Schiermonnikoog (53°30’ N, 6°10’ E), the Netherlands (Bakker 1989). The average annual temperature is 9 °C, and the average annual rainfall is 807 mm (data from www.knmi.nl). In this ecosystem, the western part (the study area) of the salt marsh has undergone more than 100 years’ succession (Olff, De Leeuw, Bakker, Platerink, & Van Wijnen, 1997; Fig. 1) and is dominated by the tall late successional grass, *Elytrigia atherica*, when cattle grazing is absent. Primary productivity is high (1119.8 ± 201.4 g dw m^−2^ y^−1^; mean ± 1 se, n= 4; measured in 2018) in this area.

**Fig. 1.**
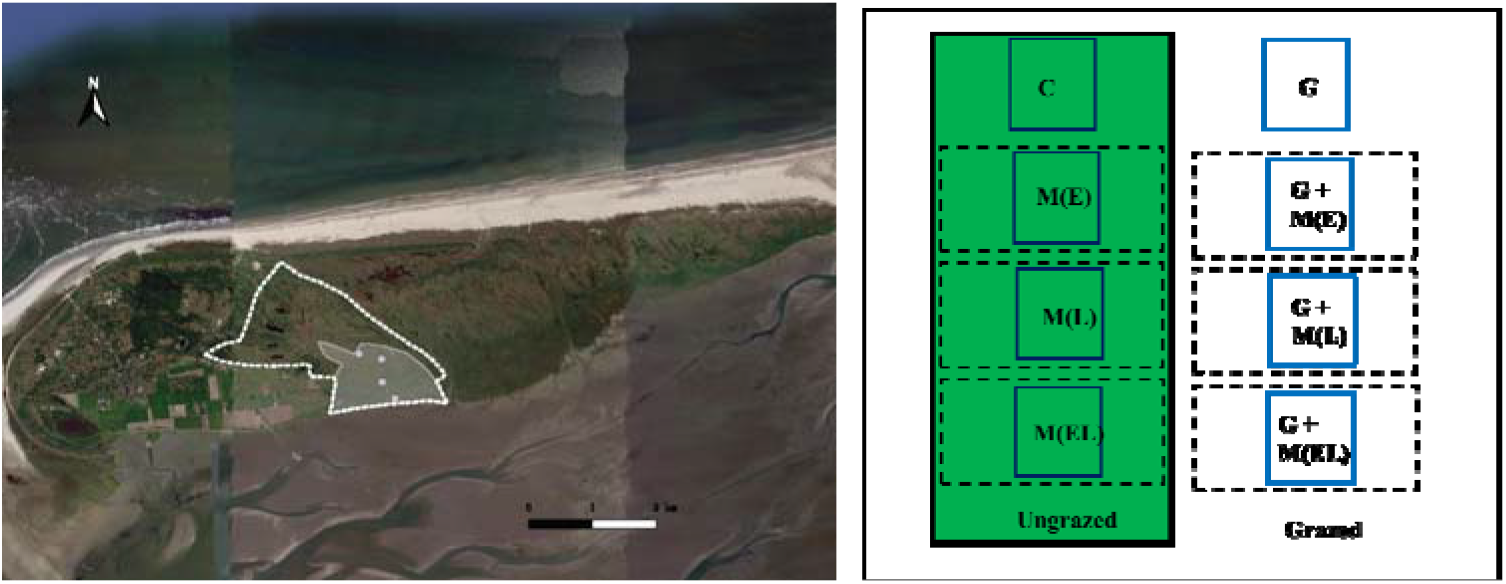
Description of the study site (left panel) and experimental design (right panel). The drawn grey area has been under cattle grazing since 1972, grazing area expanded to the dotted white area since 1993. The four white dots represent 4 blocks. Treatments for one block are shown in the right panel. Dashed black rectangles were subjected to different mowing treatments, permanent plots (blue rectangles) were established within treatments. In the field, treatments were randomized within blocks. The size of the exclosures and permanent plots are not projected according to their actual measurements. C: control, i.e. land abandonment without grazing and mowing; G: cattle grazing; M (E): early season mowing; M (L): late season mowing; M (EL): both early and late season mowing; G + M (E): cattle grazing plus early season mowing; G + M (L): cattle grazing plus late season mowing; G + M (EL): cattle grazing plus both early and late season mowing.

### Grazing history

In the 1950s, part of the study area was cut for hay, and part was grazed by heifers from farmers of the island and mainland. The animals from the mainland came to the island by boat in spring and left in autumn. This transport stopped in 1958 as the heifers often got affected by liver fluke. Later, grazing was restricted to a small part of the salt marsh, while the remaining area was abandoned. Subsequently, plant community in the abandoned area became dominated by *E. atherica*, and plant diversity declined over the following 10 years (Bakker 1989). As local farmers got more heifers and intended to extend the grazed area in the salt marsh, and conservation managers wanted to reverse the trend of biodiversity decline, cattle grazing with heifers restarted in 1972. Cattle graze from May to November in this area, after which they were taken out by farmers and moved indoors. Stocking density reduced from 1.5 to 0.5 heads ha^−1^ from 1993 onwards, as the potential area that could be grazed expanded (Bakker, De Vlas, & Van Tooren, 1993; Fig. 1).

### Experimental design

The second author (JPB) established four experimental blocks to monitor the effects of cattle grazing, mowing, and their combinations on plant communities in 1972. Although mowing became rare in salt marshes in recent decades (Esselink, 2017), mowing and grazing plus mowing were included in the experiment because these management tools are common in semi-natural grasslands in Europe (Tälle et al., 2016). Each block contains grazed and ungrazed (exclosure) parts. Exclosures (ca. 8 m × 42 m) were constructed with two electrical strands running 0.5 and 1 m above the ground. Each block contained eight treatments, including 1) an undisturbed control (C, i.e. land abandonment; without grazing and mowing), 2) cattle grazing (G), 3) early season mowing (M (E)), 4) late season mowing (M (L)), 5) both early and late season mowing (M (EL)), 6) grazing plus early season mowing (G + M (E)), 7) grazing plus late season mowing (G + M (L)), 8) grazing plus both early and late season mowing (G + M (EL)) (plot size ca. 3 m × 6 m for mowing and grazing plus mowing treatments; experimental design in Fig.1). Treatments within blocks were randomized in the field. We usually mowed in late June or early July for the early season mowing, and in late August or early September for the late season mowing. We cut the vegetation to 2 cm above ground using a brush cutter (see Fig. S1 for vegetation height in all treatments in 2018). Plant material (including litter) was raked and removed from the plots.

One permanent plot (2 m × 2 m) for each treatment was established in 1972. We recorded species occurrence and abundance in the permanent plots before the start of mowing from 1972 to 2019. Plant species occurrence and abundance were recorded by the same skilled field assistant for most years. The abundance (percent cover) was estimated using the decimal scale of Londo (1976). As the percent cover of each species was estimated independently, the total cover of living plants can sometimes exceed 100% for the multilayer canopies. We used the data from 1972, 1974-1980, 1984-1989, 2003, 2015, 2017, and 2019, as plant communities in all treatments were recorded in those years (thus 18 years of surveys). A total of 56 plant species were recorded during the 48-year experiment (see Table S1 for the full list of plant species).

### Data analysis

#### Plant diversity at the local and landscape scales

Plant diversity in the local communities was counted as the number of species in the permanent plots. Plant diversity at the landscape scale was counted as all the plant species that occurred in the four permanent plots in each treatment.

To explore whether treatments promoted particular functional groups, we classified plant species into forbs, graminoids, legumes, and woody species according to their life forms. In total, we recorded 32 forbs, 17 graminoids, 3 legumes, and 4 woody species. Legumes and woody species were relatively rare, so we do not present their results here. Additionally, we explored whether treatments impacted salt-marsh specialists (halophytes), being interesting for conservation of salt marshes. We classified halophytes according to Scherfose (1990) and Bakker et al. (2002). We recorded 17 halophytes (Table S1). We looked at treatment effects on plant diversity, diversity in different functional groups by comparing their temporal trends using generalized additive mixed models (gamm) from the R package “mgcv” (Wood 2017). See “model fitting” section for model description. Light availability and dominance are the potential explanations for changes in plant diversity (Borer et al., 2014; Koerner et al., 2018). We explored how treatments impact vegetation height, aboveground biomass, and light availability in 2018. We also explored dominance and the most dominant plants over time in each treatment (details see supplementary text; Fig. S1; Fig. S2).

#### Change in community composition in time and space

We explored whether and how treatments alter local community composition and whether they cause divergent successional trajectories. We used square root transformed cover data (to reduce the weight of the most abundant species) to construct a dissimilarity matrix using Bray–Curtis dissimilarity index. We then projected these dissimilarities in two-dimensional space using nonmetric multidimensional scaling (NMDS). To improve visualization and for a balance of temporal extent, we selected data from 1972, 1980, 1989, 2003, 2013, and 2019 (eliminating data from 1974-1979, 1984-1988, 2017).

In addition, we quantified whether and how treatments alter interannual changes in community composition and the relative importance of balanced variation and abundance gradient in these changes. Balanced variation reflects changes in relative abundances among species with total abundance unchanged. In contrast, abundance gradient indicates changes in total abundance with relative abundances of species unchanged (Baselga, 2013). In other words, higher values of the abundance gradient indicate higher species loss/gain (in abundances) while higher values of the balanced variation indicate higher species replacement. We calculated temporal community dissimilarity for a local community (permanent plot) in a given time point compared with the previous time point (e.g. 1974 to 1972, 2019 to 2017) based on cover data. We used the function “beta.pair.abund” from the R package “betapart” (Baselga & Orme, 2012) with the Bray–Curtis dissimilarity index to calculate total dissimilarity and partition it into balanced variation and abundance gradient.

Further, we looked at whether treatments cause vegetation homogenization in the landscape. Specifically, we evaluated the overall dissimilarity across four permanent plots in each treatment based on cover data. We used function “beta.multi.abund” from the R package “betapart” (Baselga, 2017; Baselga & Orme, 2012) with the Bray–Curtis dissimilarity index to calculate total dissimilarity and partition it into balanced variation and abundance gradient.

#### Model fitting

We fitted long-term trends for plant diversity, diversity in different functional groups at the local and landscape scales, and temporal and spatial community dissimilarity. We fitted these trends using generalized additive mixed models (gamm) from the R package “mgcv” (Wood 2017). For models fitted in the local communities, treatment was the fixed variable, smooths fitted for each treatment, the block was a random variable. Temporal autocorrelation was adjusted using the corCAR1 model. The autocorrelation structure was not retained if its inclusion did not significantly improve model fit (ΔAIC < 4; details see Table S2-4). We fitted similar models at the landscape scale except that the random effect of the block was removed because we obtained one value per treatment (i.e. across blocks; see supplementary text for model specification). As plant diversity varied in different treatments at the start of the experiment, here we focused on comparing the trends in different treatments. We estimated the trends using restricted maximum likelihood (REML). Trends are significant when p < 0.05. Data were analyzed in R 4.0.2 (R Core Team, 2020).

## Results

### Plant diversity at the local and landscape scales

Over time, plant diversity declined under land abandonment (without grazing and mowing) and late season mowing both at the local and landscape scales. Plant diversity decline under land abandonment was more apparent at the landscape scale: 48 years after the start of the experiment, plant diversity declined by 4 plant species on average in the local communities and 11 species in the landscape. Conversely, plant diversity increased in the first 10 years of the experiment then followed by a decrease and leveling off under grazing plus early season mowing treatment at both spatial scales considered. The trends for plant diversity under other treatments were not significant at both scales considered (Fig. 2; Table S2).

**Fig. 2.**
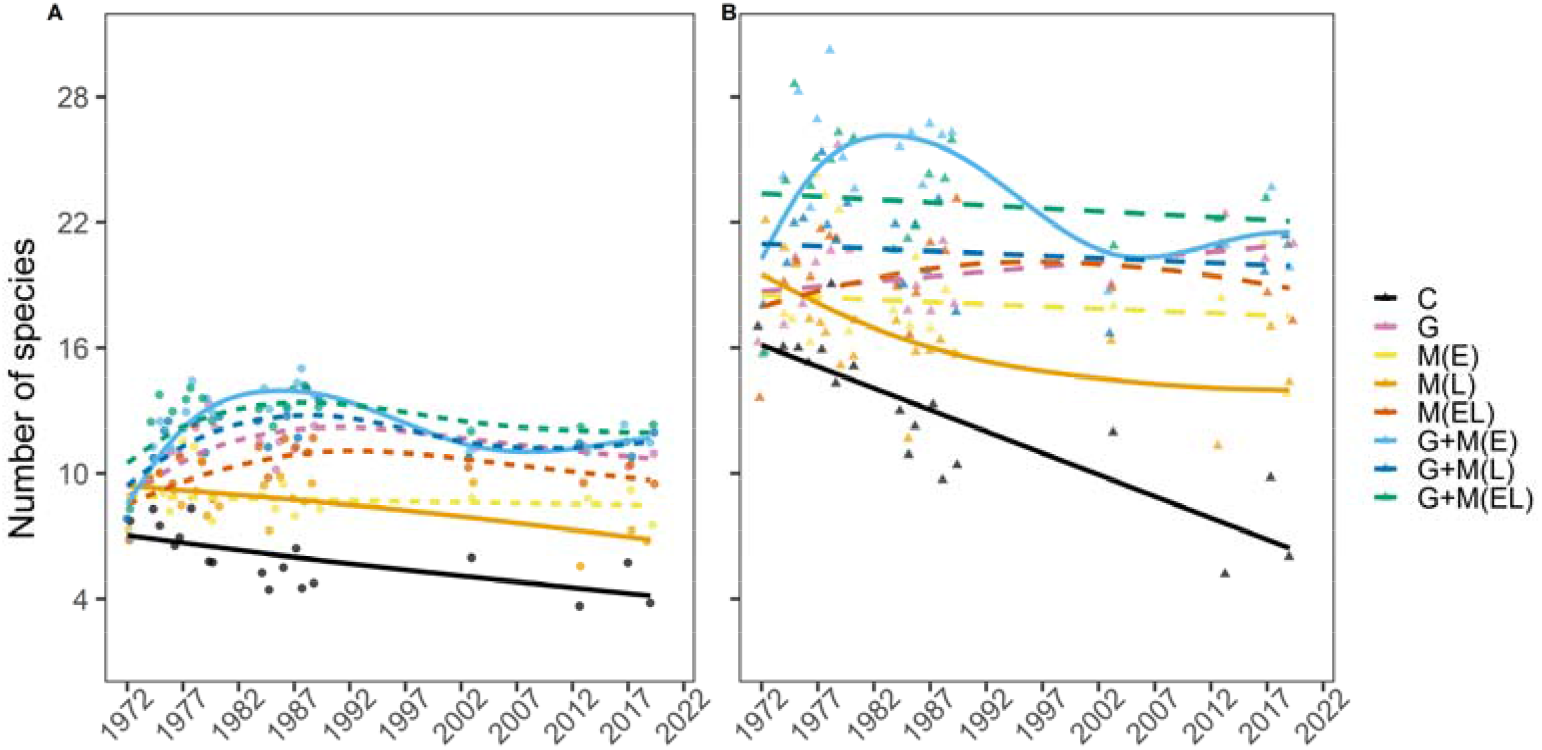
Plant diversity at the local (A) and landscape scales (B). Plant diversity was measured as the number of species. Dots are the means of the number of species over four permanent plots (2 m × 2 m). Triangles are the number of species in four permanent plots in each treatment. The solid lines represent significant trends (P < 0.05), dashed lines represent non-significant trends (p > 0.05). Lines are fitted with the generalized additive mixed model (gamm; Table S2). Treatment descriptions correspond to those of Fig. 1.

Decreased plant diversity under land abandonment was due to a decrease in graminoids at the local scale, while a decrease in graminoids and halophytes at the landscape scale. Decreased plant diversity under late season mowing at both scales was due to a decrease in graminoids. Graminoids decreased slightly (also fluctuated), while forbs increased over time at the landscape scale under grazing. Similarly, graminoids decreased slightly (also strong fluctuated), while forbs increased over time, halophytes also increased but leveled off at the end of the experiment at the landscape scale under early and late season mowing treatment. Forbs increased over time at the local scale under early season mowing. Graminoids, forbs, and halophytes fluctuated or did not change under other treatments (Fig. 3; Table S3).

**Fig. 3.**
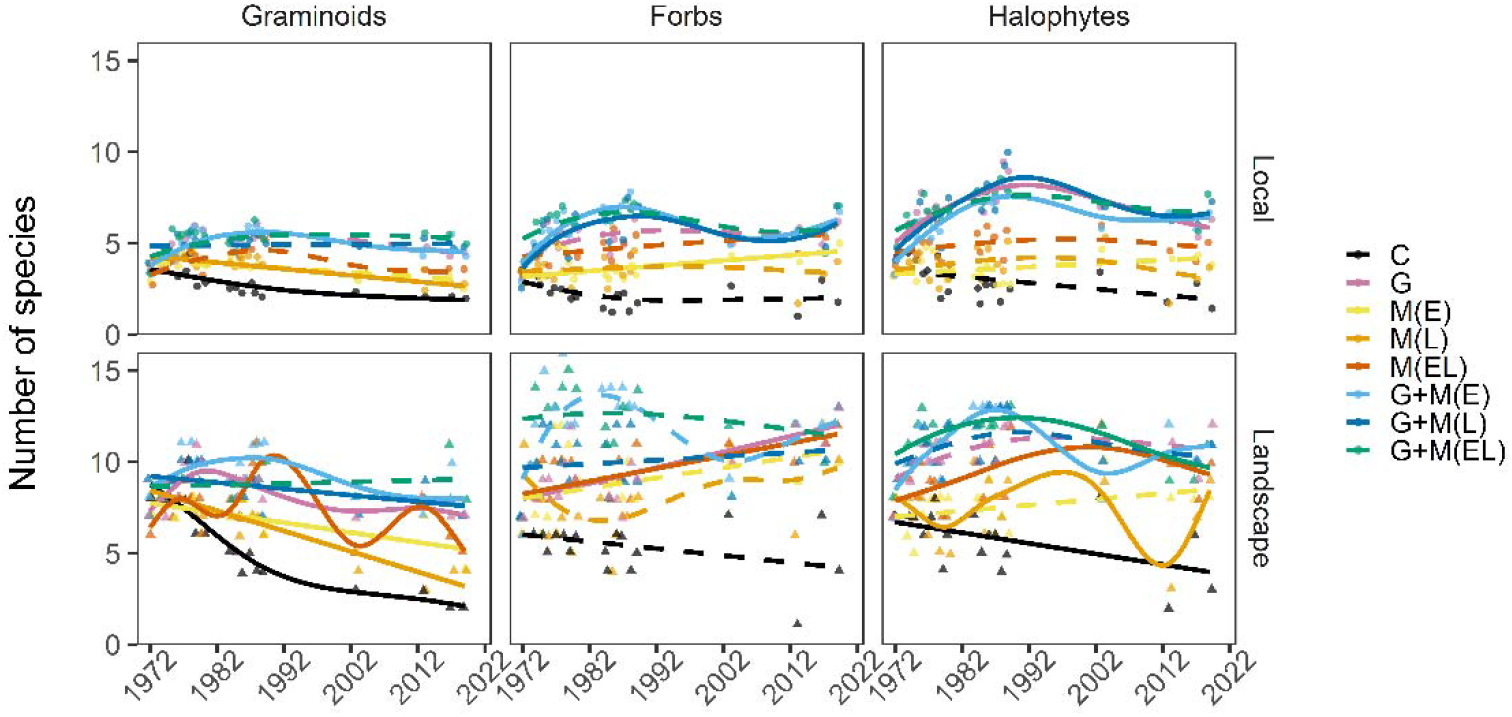
Number of graminoids, forbs, and halophytes over time at the local and landscape scales. Halophytes are plant species well adapted to saline and anoxic conditions. The solid lines represent significant trends (P < 0.05), dashed lines represent non-significant trends (p > 0.05). Lines are fitted with the generalized additive mixed model (gamm; Table S3). Treatment descriptions correspond to those of Fig. 1.

### Change in community composition in time and space

Local plant communities in all treatments underwent different successional trajectories, particularly those under land abandonment (black dots and lines in Fig. 4). Plant communities under all treatments, except for that under grazing, deviated from their initial state 48 years after the start of the experiment. Plant communities in the early season mowing and the early and late season mowing underwent similar successional trajectories (top in NMDS in Fig. 4). The successional trajectories of plant communities under grazing plus mowing treatments were generally similar to that under grazing (Fig. 4).

**Fig. 4.**
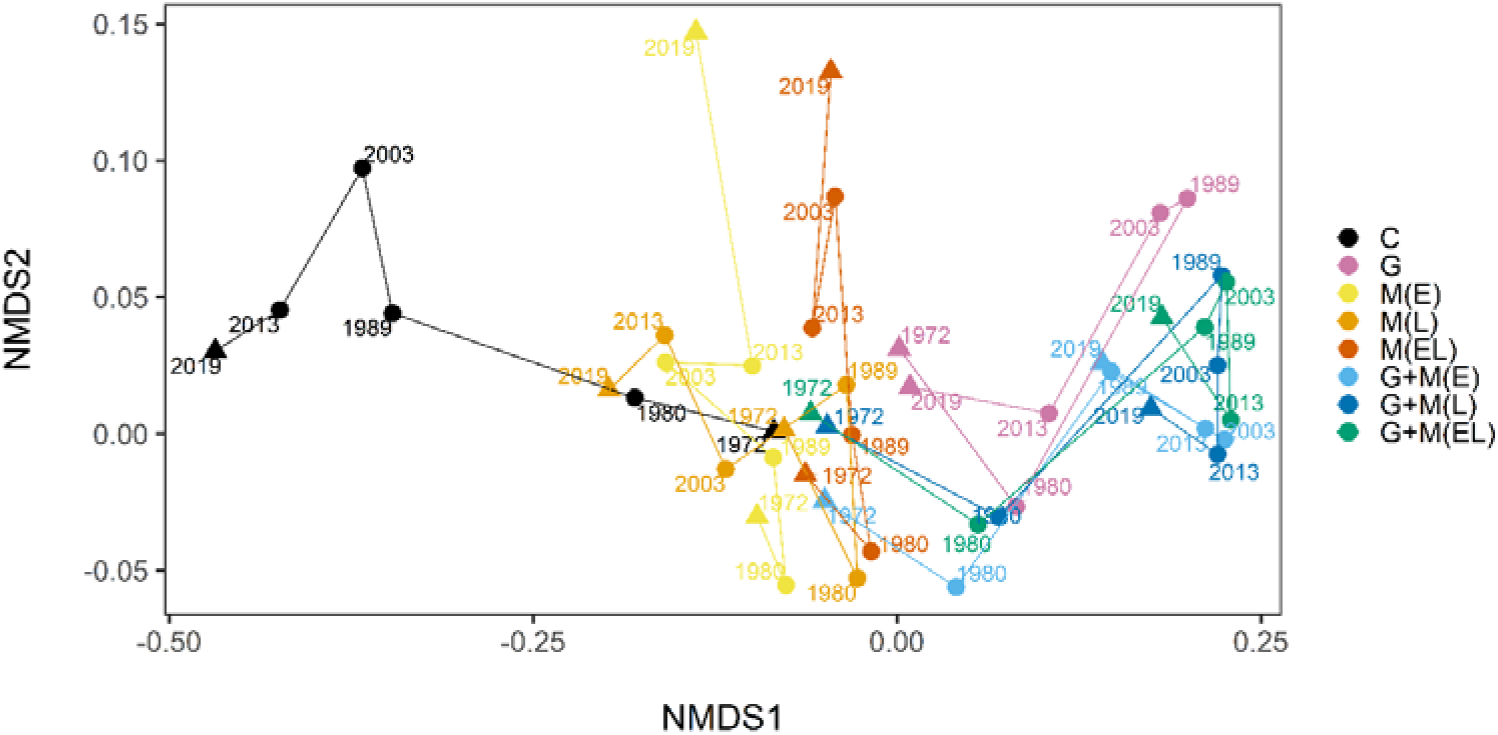
Nonmetric multidimensional scaling (NMDS) ordination plot of community dissimilarity in two-dimensional space. Community dissimilarity was based on square root transformed abundance data (to reduce the weight of the most abundant species) using the Bray–Curtis dissimilarity index. To improve visualization and for a balanced temporal extent, data from 1972, 1980, 1989, 2003, 2013, and 2019 are shown (eliminating data from 1974-1979, 1984-1988, 2017). Each point represents the community composition of a given treatment and year (averaged over four permanent plots). Triangles indicate the start and end of the experiment. The distance between any two points represents the dissimilarity of the two communities. Stress = 0.18. Treatment descriptions correspond to those of Fig. 1.

Under all treatments, temporal community dissimilarity was more attributable to balanced variation. Temporal community dissimilarity did not change over time under land abandonment and grazing plus early season mowing. Under land abandonment, balanced variation did not change over time but abundance gradient decreased slightly. Under grazing plus early season mowing, balanced variation decreased in the first 10 years but increased afterward, abundance gradient decreased slightly over time. Temporal community dissimilarity increased over time under grazing and late season mowing treatments. It decreased in the first half of the experiment but increased in the second half of the experiment under other treatments (upper panel in Fig. 5; Table S4). Spatial community dissimilarity was entirely attributable to balanced variation. Spatial community dissimilarity declined over time under land abandonment. It decreased slightly in the first half of the experiment but increased in the second half under early season mowing. It did not change under other treatments (lower panel in Fig. 5; Table S4).

**Fig. 5.**
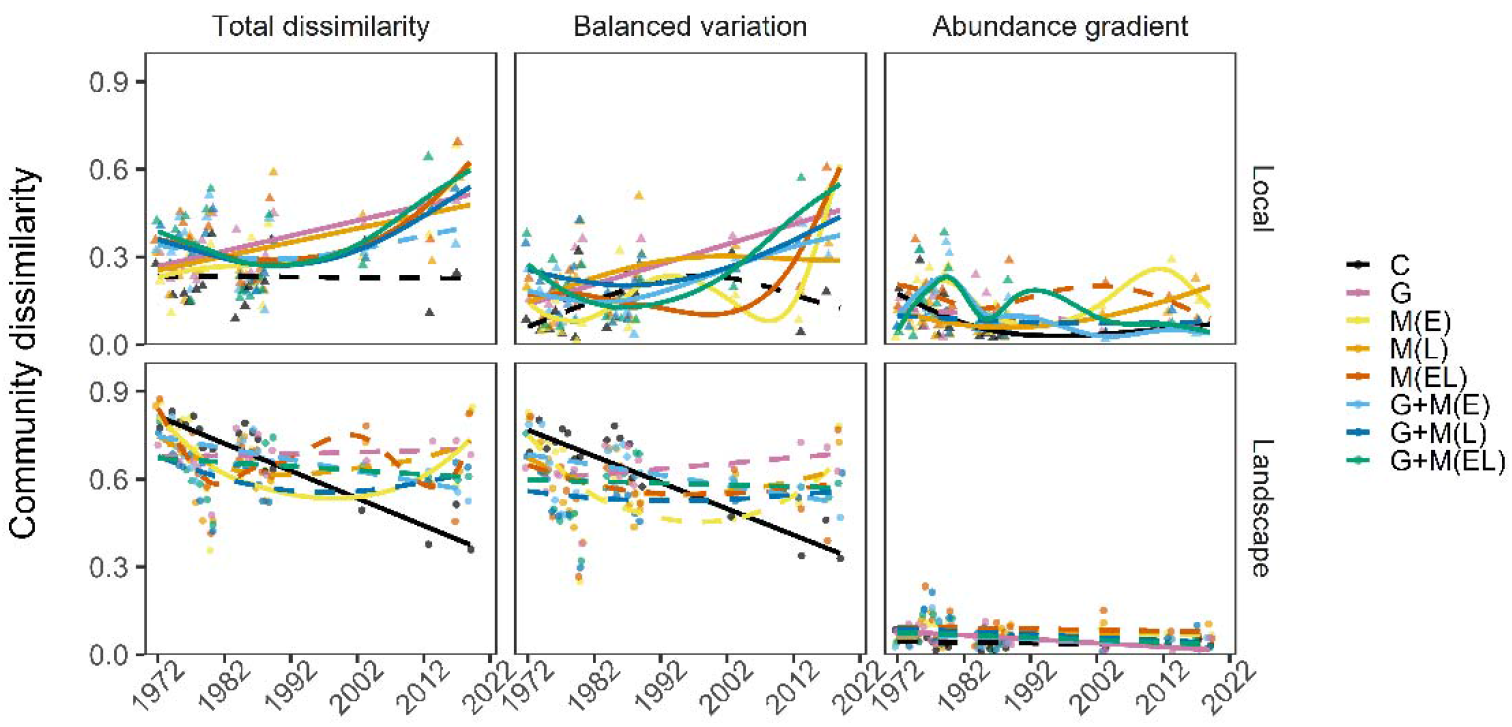
Temporal and spatial community dissimilarity. The upper panel of Fig. 5 is similar to Fig. 4 but shows the dissimilarity of a local community in a given time point compared with the previous time point. Temporal community dissimilarity was calculated as a given time point compared with the previous time point based on cover data for each permanent plot. Spatial community dissimilarity was calculated as the total dissimilarity based on cover data across four permanent plots in each treatment. Dots are the means over four permanent plots. Triangles are the dissimilarities of four permanent plots in each treatment each year. The solid lines represent significant trends (P < 0.05), dashed lines represent non-significant trends (p > 0.05). Lines are fitted with the generalized additive mixed model (gamm; Table S4). Treatment descriptions correspond to those of Fig. 1.

## Discussion

Our long-term experiment demonstrates that plant diversity and community composition in time and space under land abandonment were strongly different from that under management regimes (grazing, mowing, grazing plus mowing). Whereas, all management regimes, except the late season mowing, maintained plant diversity at both spatial scales considered. Additionally, all management regimes maintained vegetation heterogeneity in the landscape. However, local community composition under different management regimes changed differently over time.

### Plant communities without management (abandonment)

Without grazing and mowing, the late successional grass, *E. atherica*, replaced other subordinate plant species and became dominant in the local communities (Fig. S2). Subsequently, plant diversity declined. Also, rapid expansion of this grass caused vegetation homogenization over time in the landscape (Fig. 5), which worsen the decline in plant diversity at the landscape scale, particularly for graminoids and halophytes. Such an increase in *E. atherica* accompanying a decrease in plant diversity is widely observed in European salt marshes (Milotić, Erfanzadeh, Pétillon, Maelfait, & Hoffmann, 2010; Pétillon, Ysnel, Canard, & Lefeuvre, 2005; Rupprecht, Wanner, Stock, & Jensen, 2015; Veeneklaas, Dijkema, Hecker, & Bakker, 2013). Therefore, long-term management is needed to conserve plant diversity in many European salt marshes (Chen et al., 2020).

### Comparing grazing and mowing

Plant diversity and community composition were generally similar under grazing and grazing plus mowing. Although rare in salt marshes (Esselink, 2017), grazing plus mowing is still commonly used in European grasslands (Tälle et al., 2016). Our experiment shows that grazing plus mowing may be more effective in productive systems. Grazing plus mowing can suppress the dominant plant (*E. atherica*) better than grazing or mowing alone, especially for blocks/areas initially dominated by this grass (Fig. S2). Interannual composition change in local communities under all management regimes was mainly driven by replacement in species identities and abundances (upper panel, Fig. 5). It generally increased under all management regimes except for that under grazing plus early season mowing. It suggests that grazing plus early season mowing may be particularly suitable for maintaining community compositional stability in salt marshes. Contrary to what we expected, grazing did not promote graminoids while decreasing forbs. Instead, both grazing and the early and late season mowing promoted forb species, but only at the landscape scale. This may be because cattle are more selective with plant height rather than their life forms (Díaz et al., 2007). In other words, taller plants regardless of graminoids or forbs are more likely to be grazed than shorter ones. Thus, in a forb-rich system such as ours, small forbs can still germinate and establish since grazing and mowing increased light availability via removing biomass (Fig. S1) or reducing dominance (Fig. S2). Indeed, Ludvíková et al., (2014) also found that short forbs increase under mowing and herbivore trampling treatments, while graminoids are not responsive to these treatments in a Central European grassland. Surprisingly, the trends for number of halophytes did not differ much under grazing, mowing, and grazing plus mowing regimes (did not decrease or fluctuated over time). This suggests that the establishment of salt-marsh specialists is more limited by light rather than anoxic conditions (Bakker, Dijkstra, & Russchen, 1985). Therefore, grazing and mowing generally had similar effects on diversity in different functional groups. Similar to that of grazing, the early season mowing and early and late season mowing regimes maintained plant diversity. However, the successional trajectories of the local plant communities under these mowing regimes were different from that under grazing. This is particularly the case in the first half of the experiment (1972-1989) (Fig. 4). Kahmen et al., (2002) also found that local communities under grazing and mowing followed different successional trajectories, but they were generally more similar when compared with land abandonment and other management tools (e.g. burning) in a 25-year experiment in a German grassland. This difference in successional trajectories may because biomass was only removed once or twice per year under mowing regimes, while biomass was removed continuously under grazing during the growing season. This pattern may also arise because herbivore trampling and deposition of urine and dung mediates plant communities’ response via altering soil properties (Schrama et al., 2013). Indeed, we found that halophyte *Juncus gerardii* was more abundant under grazing and grazing plus mowing regimes, while glycophyte *Festuca rubra* was more abundant under mowing regimes (Fig. S2). In the field, these two species looked very similar, but *J. gerardii* may be better adapted to anoxic conditions under grazing.

### Comparing the effects of timing of mowing

Our results suggest that timing and frequency of mowing had a strong impact on plant communities, which is consistent with previous analyses (Dee, Thomas, Thompson, & Palmer, 2016; Fynn, Morris, & Edwards, 2004; Huhta, Rautio, Tuomi, & Laine, 2001; Wilson & Clark, 2001). Plant diversity declined, while successional trajectories under late season mowing deviated from that of under early and early and late season mowing regimes. As plants usually germinate in the early growing season, early season mowing is likely to provide more opportunities for species germination and establishment via increasing light availability. Indeed, the early season mowing and the early and late season mowing regimes generally had similar effects on plant diversity and community composition.

### Long-term across-scale monitoring

Our results suggest that plant communities take a long time to respond to management regimes, and such long-term experiments are valuable. We found that interannual composition change under some management regimes decreased in the first half of the experiment but increased in the second half of the experiment. Also, plant communities under grazing diverged from their initial states in the first half of the experiment but became similar again in the second half of the experiment. This may be due to reduced stocking density after 1993. The importance of long-term experiments in revealing the effects of herbivores on plant communities has also been confirmed by other studies. For instance, Porensky *et al.* (2017) found that ungrazed and lightly grazed pastures experienced relatively large shifts in plant community composition compared with that under moderate and heavy grazing, particularly in the last 25 years of a 75-year sustained yearling grazing experiment in a US shortgrass steppe. Similarly, cross-scale monitoring is important (Collins, Glenn, & Briggs, 2002). We show here that plant diversity decline was stronger at the landscape scale than at the local scale under land abandonment. Therefore, results based on shorter-term and local-scale experiments may inaccurately estimate land abandonment or management regimes on plant communities.

### Implications

Our results have clear implications for conservation and management of biodiversity in salt marshes. All management regimes tested here, except the late season mowing, can maintain plant diversity across scales in the long term. All management regimes can maintain vegetation heterogeneity in the landscape over time. As vegetation heterogeneity was mostly driven by high turnover in species identities and abundances between patches (blocks), conservation should focus on larger spatial scales or multiple sites. However, grazing and mowing may drive local communities in different successional trajectories. Also, all management regimes, except the grazing plus early season mowing, may increase interannual community composition change. Economically speaking, early season mowing may be more viable than early and late season mowing, grazing may be more viable than grazing plus mowing. There are many other factors to be considered, for example, the accessibility of livestock. Therefore, the optimal management tool in a system may be highly dependent on the goal of conservation and availability of resources.

## Supporting information

Supplementary file for Long-term cross-scale comparison of grazing and mowing on plant diversity and community composition in a salt-marsh system

## Author contributions

JPB designed and conducted the experiments. CS and QC maintained the experiment and collected data since 2012, and 2016, respectively. QC and JA conceived the idea, QC analyzed the data and wrote the manuscript. All authors contributed to revisions and gave final approval for publication.

## Acknowledgments

We thank Yzaak de Vries for recording the occurrence and abundance of plant species in permanent plots for many years. We thank Richard Ubels for helping record the occurrence and abundance of plant species. We thank Jacob Hogendorf, Jan van den Burg, and Ron Snijders for helping mowing in the field. We thank Ido Pen for his helpful suggestions for data analysis. We thank Yong-fei Bai for his constructive comments on an early draft. We thank two anonymous reviewers and the editor for their constructive comments and suggestions to improve this manuscript. We thank Ron Snijders for his constructive comments to improve the readability of this manuscript. We thank Natuurmonumenten for offering us the opportunity to work in the salt marsh of the island of Schiermonnikoog. QC is funded by CSC (China Council Scholarship). JA was supported by a Visitor’s Travel Grant (040.11.631) of the Netherlands Organisation for Scientific Research (NWO).

## Data and materials availability

data and the associated R code for producing figures will be deposited in the Dryad Digital Repository once the manuscript gets accepted.

## References

Bakker, J.P. (1989). Nature management by grazing and cutting. Kluwer Academic Publishers, Dordrecht.

Bakker, J. P., Esselink, P., Dijkema, K. S., Van Duin, W. E., & De Jong, D. J. (2002). Restoration of salt marshes in the Netherlands. Hydrobiologia, 478, 29–51. doi: 10.1023/A:1021066311728 (Original work published)

Bakker, J.P., De Vlas, J., & Van Tooren, B. F. (1993). Uitbreiding begrazing van de Oosterkwelder op Schiermonnikoog. De Levende Natuur, 94, 118–122. (Original work published)

Bakker, J P, Bos, D., & De Vries, Y. (2003). To graze or not to graze: that is the question. Challenges to the Wadden Sea - Proceedings of the 10th International Scientific Wadden Sea Symposium, 67–87. (Original work published)

Bakker, J P, Dijkstra, M., & Russchen, P. T. (1985). Dispersal, germination and early establishment of halophytes and glycophytes on a grazed and abandoned salt-marsh gradient. New Phytologist, 101(2), 291–308. doi: 10.1111/j.1469-8137.1985.tb02836.x (Original work published)

Bakker, Jan P. (1978). Changes in a salt-marsh vegetation as a result of grazing and mowing - a five-year study of permanent plots. Vegetatio, 38(2), 77–87. (Original work published)

Baselga, A. (2013). Separating the two components of abundance-based dissimilarity: Balanced changes in abundance vs. abundance gradients. Methods in Ecology and Evolution, 4(6), 552–557. doi: 10.1111/2041-210X.12029 (Original work published)

Baselga, A. (2017). Partitioning abundance-based multiple-site dissimilarity into components: balanced variation in abundance and abundance gradients. Methods in Ecology and Evolution, 8(7), 799–808. doi: 10.1111/2041-210X.12693 (Original work published)

Baselga, A., & Orme, C. D. L. (2012). Betapart: An R package for the study of beta diversity. Methods in Ecology and Evolution, 3(5), 808–812. doi: 10.1111/j.2041-210X.2012.00224.x (Original work published)

Bignal, E. M., & McCracken, D. I. (1996). Low-Intensity Farming Systems in the Conservation of the Countryside. Journal of Applied Ecology, 33(3), 413–424. (Original work published)

Borer, E. T., Seabloom, E. W., Gruner, D. S., Harpole, W. S., Hillebrand, H., Lind, E. M., … Yang, L. H. (2014). Herbivores and nutrients control grassland plant diversity via light limitation. Nature, 508(7497), 517–520. doi: 10.1038/nature13144 (Original work published)

Briske, D.D. & Richards, J.H. (1995) Plant responses to defoliation: a physiological, morphological and demographic evaluation. Wildland Plants: Physiological Ecology and Developmental Morphology (eds D.J. Bedunah & R.E. Sosebee), pp. 635–710. Society for Range Management, Denver, CO, USA.

Catorci, A., Cesaretti, S., Malatesta, L., & Tardella, F. M. (2014). Effects of grazing vs mowing on the functional diversity of sub-Mediterranean productive grasslands. Applied Vegetation Science, 17(4), 658–669. doi: 10.1111/avsc.12103 (Original work published)

Chase, J. M., McGill, B. J., McGlinn, D. J., May, F., Blowes, S. A., Xiao, X., … Gotelli, N. J. (2018). Embracing scale-dependence to achieve a deeper understanding of biodiversity and its change across communities. Ecology Letters, 1737–1751. doi: 10.1111/ele.13151 (Original work published)

Chen, Q., Bakker, J. P., Alberti, J., & Smit, C. (2020). Long-term management is needed for conserving plant diversity in a Wadden Sea salt marsh. Biodiversity and Conservation, 29(7), 2329–2341. doi: 10.1007/s10531-020-01976-w (Original work published)

Collins, S. L., Glenn, S. M., & Briggs, J. M. (2002). Effect of local and regional processes on plant species richness in tallgrass prairie. Oikos, 99(3), 571–579. doi: 10.1034/j.1600-0706.2002.12112.x (Original work published)

Collins, S. L., Knapp, A. K., Briggs, J. M., Blair, J. M., & Steinauer, E. M. (1998). Modulation of diversity by grazing and mowing in native tallgrass prairie. Science, 280(5364), 745–747. doi: 10.1126/science.280.5364.745 (Original work published)

Davidson, K. E., Fowler, M. S., Skov, M. W., Doerr, S. H., Beaumont, N., & Griffin, J. N. (2017). Livestock grazing alters multiple ecosystem properties and services in salt marshes: a meta-analysis. Journal of Applied Ecology, 54(5), 1395–1405. doi: 10.1111/1365-2664.12892 (Original work published)

Davies, K. F., Chesson, P., Harrison, S., Inouye, B. D., Melbourne, B. A., & Rice, K. J. (2005). Spatial Heterogeneity Explains the Scale Dependence of the Native-Exotic Diversity Relationship. Ecology, 86(6), 1602–1610. (Original work published)

De Cauwer, B., & Reheul, D. (2009). Impact of land use on vegetation composition, diversity and potentially invasive, nitrophilous clonal species in a wetland region in Flanders. Agronomy for Sustainable Development, 29(2), 277–285. doi: 10.1051/agro:2008049 (Original work published)

Dee, J. R., Thomas, S. M., Thompson, S. D., & Palmer, M. W. (2016). Long-term late season mowing maintains diversity in southern US tallgrass prairie invaded by Bothriochloa ischaemum. Applied Vegetation Science, 19(3), 442–453. doi: 10.1111/avsc.12227 (Original work published)

Díaz, S., Lavorel, S., McIntyre, S., Falczuk, V., Casanoves, F., Milchunas, D. G., … Campbell, B. D. (2007). Plant trait responses to grazing - A global synthesis. Global Change Biology, 13(2), 313–341. doi: 10.1111/j.1365-2486.2006.01288.x (Original work published)

Elschot, K., Bakker, J. P., Temmerman, S., Van De Koppel, J., & Bouma, T. J. (2015). Ecosystem engineering by large grazers enhances carbon stocks in a tidal salt marsh. Marine Ecology Progress Series, 537, 9–21. doi: 10.3354/meps11447 (Original work published)

Esselink, P. (2017). Wadden Sea Quality Status Report. Wadden Sea Ecosystem, 9. Retrieved from http://cwss.www.de/TMAP/Qsr99/Qsr99.html (Original work published)

Fynn, R. W. S., Morris, C. D., & Edwards, T. J. (2004). Effect of burning and mowing on grass and forb diversity in a long-term grassland experiment. Applied Vegetation Science, 7(1), 1–10. doi: 10.1111/j.1654-109X.2004.tb00589.x (Original work published)

Garrido, P., Mårell, A., Öckinger, E., Skarin, A., Jansson, A., & Thulin, C. G. (2019). Experimental rewilding enhances grassland functional composition and pollinator habitat use. Journal of Applied Ecology, 56(4), 946–955. doi: 10.1111/1365-2664.13338 (Original work published)

Gedan, K. B., Silliman, B. R., & Bertness, M. D. (2009). Centuries of Human-Driven Change in Salt Marsh Ecosystems. Annual Review of Marine Science, 1(1), 117–141. doi: 10.1146/annurev.marine.010908.163930 (Original work published)

Gillet, F., Kohler, F., Vandenberghe, C., & Buttler, A. (2010). Effect of dung deposition on small-scale patch structure and seasonal vegetation dynamics in mountain pastures. Agriculture, Ecosystems and Environment, 135(1-2), 34–41. doi: 10.1016/j.agee.2009.08.006 (Original work published)

Huhta, A.-P., Rautio, P., Tuomi, J., & Laine, K. (2001). Restorative mowing on an abandoned semi-natural meadow: short-term and predicted long-term effects. Journal of Vegetation Science, 12(5), 677–686. doi: 10.2307/3236908 (Original work published)

Isbell, F., Gonzalez, A., Loreau, M., Cowles, J., Díaz, S., Hector, A., … Larigauderie, A. (2017). Linking the influence and dependence of people on biodiversity across scales. Nature, 546(7656), 65–72. doi: 10.1038/nature22899 (Original work published)

Kahmen, S., Poschlod, P., & Schreiber, K. F. (2002). Conservation management of calcareous grasslands. Changes in plant species composition and response of functional traits during 25 years. Biological Conservation, 104(3), 319–328. doi: 10.1016/S0006-3207(01)00197-5 (Original work published)

Kobayashi, T., Hori, Y., & Nomoto, N. (1997). Effects of trampling and vegetation removal on species diversity and micro-environment under different shade conditions. Journal of Vegetation Science, 8(6), 873–880. doi: 10.2307/3237032 (Original work published)

Koerner, S. E., Smith, M. D., Burkepile, D. E., Hanan, N. P., Avolio, M. L., Collins, S. L., … Zelikova, T. J. (2018). Change in dominance determines herbivore effects on plant biodiversity. Nature Ecology & Evolution, 2, 1925–1932. doi: 10.1038/s41559-018-0696-y (Original work published)

Kohler, F., Gillet, F., Gobat, J. M., & Buttler, A. (2004). Seasonal vegetation changes in mountain pastures due to simulated effects of cattle grazing. Journal of Vegetation Science, 15(2), 143–150. doi: 10.1111/j.1654-1103.2004.tb02249.x (Original work published)

Kumm, K. I. (2003). Sustainable management of Swedish seminatural pastures with high species diversity. Journal for Nature Conservation, 11(2), 117–125. doi: 10.1078/1617-1381-00039 (Original work published)

Lepš, J. (2014). Scale- and time-dependent effects of fertilization, mowing and dominant removal on a grassland community during a 15-year experiment. Journal of Applied Ecology, 51, 978–987. doi: 10.1111/1365-2664.12255 (Original work published)

Lezama, F., & Paruelo, J. M. (2016). Disentangling grazing effects: trampling, defoliation and urine deposition. Applied Vegetation Science, 19(4), 557–566. doi: 10.1111/avsc.12250 (Original work published)

Londo, G. (1976). The decimal scale for releves of permanent quadrats. Vegetatio, 33(1), 61–64. (Original work published)

Ludvíková, V., Pavlů, V. V., Gaisler, J., Hejcman, M., & Pavlů, L. (2014). Long term defoliation by cattle grazing with and without trampling differently affects soil penetration resistance and plant species composition in Agrostis capillaris grassland. Agriculture, Ecosystems and Environment, 197, 204–211. doi: 10.1016/j.agee.2014.07.017 (Original work published)

Lundholm, J. T., & Larson, D. W. (2003). Relationships between Spatial Environmental Heterogeneity and Plant Species Diversity on a Limestone Pavement. Ecography, 26(6), 715–722. (Original work published)

Mikola, J., Setälä, H., Virkajärvi, P., Saarijärvi, K., Ilmarinen, K., Voigt, W., & Vestberg, M. (2009). Defoliation and patchy nutrient return drive grazing effects on plant and soil properties in a dairy cow pasture. Ecological Monographs, 79(2), 221–244. doi: 10.1890/08-1846.1 (Original work published)

Milotić, T., Erfanzadeh, R., Pétillon, J., Maelfait, J. P., & Hoffmann, M. (2010). Short-term impact of grazing by sheep on vegetation dynamics in a newly created salt-marsh site. Grass and Forage Science, 65(1), 121–132. doi: 10.1111/j.1365-2494.2009.00725.x (Original work published)

Moog, D., Poschlod, P., Kahmen, S., Schreiber, K. F. K.-F., Medicine, P., Mood, D., … Medicine, P. (2002). Comparison of species composition between different grassland management treatments after 25 years. Applied Vegetation Science, 5(1), 99–106. doi: 10.1111/j.1654-109X.2002.tb00539.x (Original work published)

Oldén, A., & Halme, P. (2016). Grazers increase β-diversity of vascular plants and bryophytes in wood-pastures. Journal of Vegetation Science, 27(6), 1084–1093. doi: 10.1111/jvs.12436 (Original work published)

Olff, H., De Leeuw, J., Bakker, J. P., Platerink, R. J., & Van Wijnen, H. J. (1997). Vegetation succesion and herbivory in a salt marsh: changes induced by sea level rise and silt deposition along an elevation gradient. Journal of Ecology, 85(6), 799–814. doi: 10.2307/2960603 (Original work published)

Olff, H., & Ritchie, M. E. (1998). Effects of herbivores on grassland plant diversity. Trends in Ecology and Evolution, 13(7), 261–265. doi: 10.1016/S0169-5347(98)01364-0 (Original work published)

Pétillon, J., Ysnel, F., Canard, A., & Lefeuvre, J. C. (2005). Impact of an invasive plant (Elymus athericus) on the conservation value of tidal salt marshes in western France and implications for management: Responses of spider populations. Biological Conservation, 126(1), 103–117. doi: 10.1016/j.biocon.2005.05.003 (Original work published)

Porensky, L. M., Derner, J. D., Augustine, D. J., & Milchunas, D. G. (2017). Plant Community Composition after 75 Yr of Sustained Grazing Intensity Treatments in Shortgrass Steppe. Rangeland Ecology and Management, 70(4), 456–464. doi: 10.1016/j.rama.2016.12.001 (Original work published)

Poschlod, P., Bakker, J. P., & Kahmen, S. (2005). Changing land use and its impact on biodiversity. Basic and Applied Ecology, 6(2), 93–98. doi: 10.1016/j.baae.2004.12.001 (Original work published)

R Core Team (2020) R: a language and environment for statistical computing. Vienna, Austria: R foundation for statistical computing. Retrieved from https://www.R-project.org/

Rupprecht, F., Wanner, A., Stock, M., & Jensen, K. (2015). Succession in salt marshes - large-scale and long-term patterns after abandonment of grazing and drainage. Applied Vegetation Science, 18(1), 86–98. doi: 10.1111/avsc.12126 (Original work published)

Schrama, M., Heijning, P., Bakker, J. P., Van Wijnen, H. J., Berg, M. P., & Olff, H. (2013). Herbivore trampling as an alternative pathway for explaining differences in nitrogen mineralization in moist grasslands. Oecologia, 172(1), 231–243. doi: 10.1007/s00442-012-2484-8 (Original work published)

Tälle, M., Deák, B., Poschlod, P., Valkó, O., Westerberg, L., & Milberg, P. (2016). Grazing vs. mowing: A meta-analysis of biodiversity benefits for grassland management. Agriculture, Ecosystems and Environment, 222, 200–212. doi: 10.1016/j.agee.2016.02.008 (Original work published)

Terres, J. M., Scacchiafichi, L. N., Wania, A., Ambar, M., Anguiano, E., Buckwell, A., … Zobena, A. (2015). Farmland abandonment in Europe: Identification of drivers and indicators, and development of a composite indicator of risk. Land Use Policy, 49, 20–34. doi: 10.1016/j.landusepol.2015.06.009 (Original work published)

Ustaoglu, E., & Collier, M. J. (2018). Farmland abandonment in Europe: an overview of drivers, consequences, and assessment of the sustainability implications. Environmental Reviews, 416(July), 396–416. (Original work published)

Van Der Graaf, A. J., Stahl, J., & Bakker, J. P. (2005). Compensatory growth of Festuca rubra after grazing: Can migratory herbivores increase their own harvest during staging? Functional Ecology, 19(6), 961–969. doi: 10.1111/j.1365-2435.2005.01056.x (Original work published)

Van Klink, R., Schrama, M., Nolte, S., Bakker, J. P., WallisDeVries, M. F., & Berg, M. P. (2015). Defoliation and Soil Compaction Jointly Drive Large-Herbivore Grazing Effects on Plants and Soil Arthropods on Clay Soil. Ecosystems, 18(4), 671–685. doi: 10.1007/s10021-015-9855-z (Original work published)

Van Klink, Roel, Nolte, S., Mandema, F. S., Lagendijk, D. D. G., WallisDeVries, M. F., Bakker, J. P., … Smit, C. (2016). Effects of grazing management on biodiversity across trophic levels–The importance of livestock species and stocking density in salt marshes. Agriculture, Ecosystems and Environment, 235, 329–339. doi: 10.1016/j.agee.2016.11.001 (Original work published)

Veeneklaas, R. M., Dijkema, K. S., Hecker, N., & Bakker, J. P. (2013). Spatio-temporal dynamics of the invasive plant species Elytrigia atherica on natural salt marshes. Applied Vegetation Science, 16(2), 205–216. doi: 10.1111/j.1654-109X.2012.01228.x (Original work published)

Wanner, A., Suchrow, S., Kiehl, K., Meyer, W., Pohlmann, N., Stock, M., & Jensen, K. (2014). Scale matters: Impact of management regime on plant species richness and vegetation type diversity in Wadden Sea salt marshes. Agriculture, Ecosystems and Environment, 182, 69–79. doi: 10.1016/j.agee.2013.08.014 (Original work published)

Wellstein, C., Otte, A., & Waldhardt, R. (2007). Seed Bank Diversity in Mesic Grasslands in Relation to Vegetation Type, Management and Site Conditions. Journal of Vegetation Science, 18(2), 153–162. (Original work published)

Wilson, M. V., & Clark, D. L. (2001). Controlling invasive Arrhenatherum elatius and promoting native prairie grasses through mowing. Applied Vegetation Science, 4(1), 129–138. doi: 10.1111/j.1654-109X.2001.tb00243.x (Original work published)

Wood, S.N. (2017) Generalized Additive Models: An Introduction with R (2^nd^ edition). Chapman and Hall/CRC.

