## Supplementary file for Long-term cross-scale comparison of grazing and mowing on plant diversity and community composition in a salt-marsh system for "Long-term cross-scale comparison of grazing and mowing on plant diversity and community composition in a salt-marsh system"

Supplementary Text

*Vegetation height, biomass, and light availability in 2018*

Vegetation height was measured by dropping a Styrofoam disc (19 cm Ø, 20 g) along a calibrated stick (n = 4 per permanent plot) in 2018. In September 2018, before the late season mowing, we measured aboveground biomass for all treatments. We clipped vegetation of two randomly chosen strips (10 cm × 100 cm) to the ground level (ca. 1 cm) adjacent to the permanent plots and weighed the biomass to the nearest 0.01 g after drying in the oven (70 °C) to constant weight. The biomass from two strips per permanent plot was added up and multiplied by 5 to estimate the g dw m^-2^ for all treatments (data presented in Fig. S2B). We also explored the underlying mechanism of changes in plant diversity via increasing light availability ( Borer *et al.* 2014). In September 2018, before the late season mowing, we measured the proportion of photosynthetically active radiation (PAR, μmole photons m^-2^ s^-1^) on a sunny day (between 12:00 and 14:00, approximately solar noon) using a light sensor (Skye, UK). We took four measurements per permanent plot. For each measurement, we simultaneously measured PAR at ground level (ca. 3.8 cm) and above vegetation (ca. 50 -100 cm). We calculated light availability as the PAR reaching the ground level compared with that of the above vegetation. Four measurements were averaged for each permanent plot. We fitted linear mixed effect models from the R package “lmerTest” (Kuznetsova *et al.* 2018) for vegetation height, aboveground biomass, and light availability, respectively. In the models, treatment was the fixed variable, the block was the random variable. Significance of fixed terms was assessed using the function ANOVA (type III), where degrees of freedom were calculated by Satterthwaite's approximation. Treatment significantly affected vegetation height (F_7,21_ = 14.20, p < 0.0001), aboveground biomass (F_7,21_ = 11.42, p < 0.0001) and light availability (F_7,21_ = 9.18, p < 0.0001). Vegetation height, aboveground biomass, and light availability was log, log, and square root transformed before fitting the models.

*Dominance, Elytrigia atherica, Festuca rubra, and Juncus gerardii*

We explored dominance because it is one of the potential explanations for the change in plant diversity (Koerner et al. 2018). Dominance was measured as the Berger-Parker dominance index, i.e. the proportional abundance of the most abundant species. We calculated the Berger-Parker dominance index using the R package “diverse” (Guevara *et al.* 2017). Additionally, we explored the percent cover of *E. atherica*, *F. rubra*, and *Juncus gerardii*, which were the most dominant plant species in the control, mowing, and grazing (as well as grazing plus mowing treatments) at the end of the experiment, respectively.

*Examples of model specification*

Models for plant diversity at the local and landscape scales were defined in R as: gamm (number of species in the local communities ~treatment + s(year, by=treatment), method="REML", random=list(block = ~1), correlation=corCAR1(form = ~ year|permanent plots)); and gamm (number of species in the landscape ~treatment + s(year, by=treatment), method="REML", correlation=corCAR1(form = ~year|treatment)), respectively.

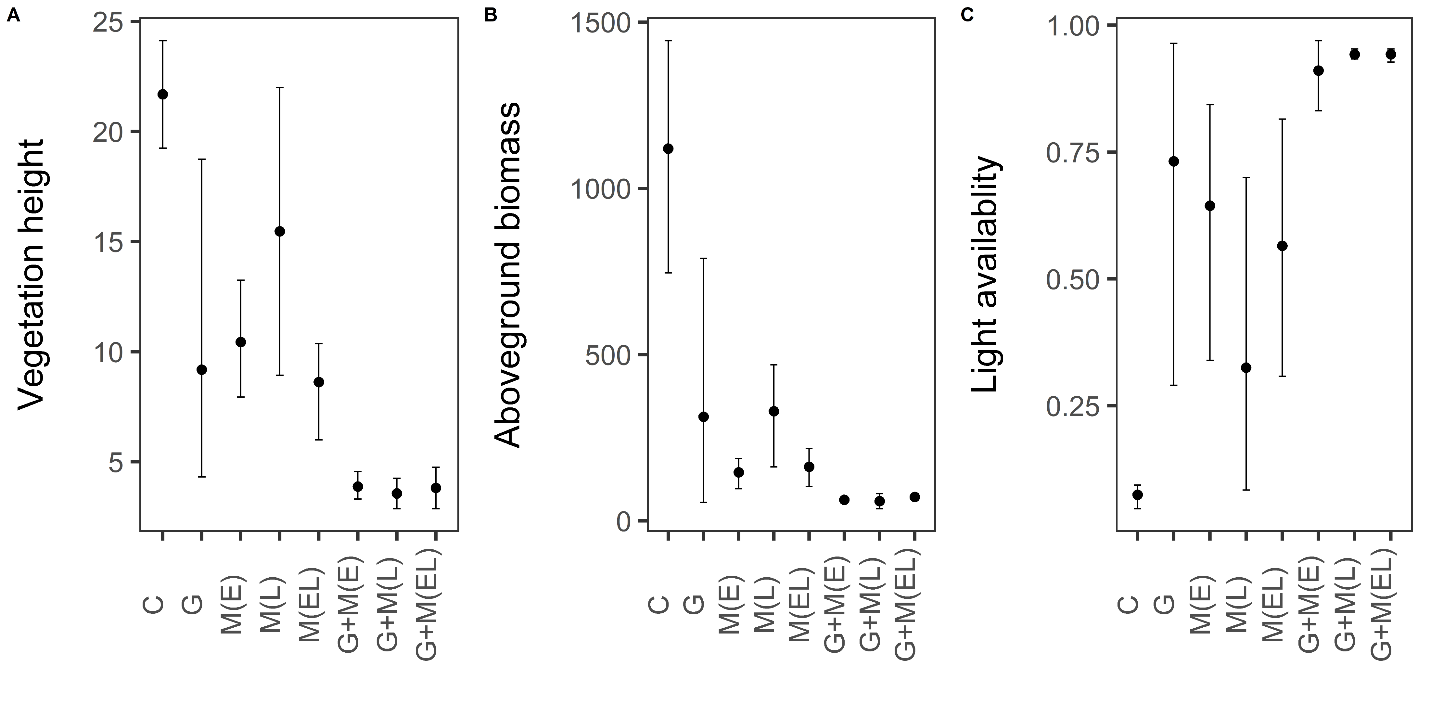
**Fig. S1. Vegetation height (cm; A), aboveground biomass (g/m**^2^**; B) and light availability (%; C) in eight treatments in 2018.** Dots represent the means of four permanent plots, error bars show the 95 % bootstrapped confidence interval using the function “mean_cl_boot” embedded in package “ggplot2”. C: control, i.e. land abandonment without grazing and mowing; G: cattle grazing; M (E): early season mowing; M (L): late season mowing; M (EL): both early and late season mowing; G + M (E): cattle grazing plus early season mowing; G + M (L): cattle grazing plus late season mowing; G + M (EL): cattle grazing plus both early and late season mowing.

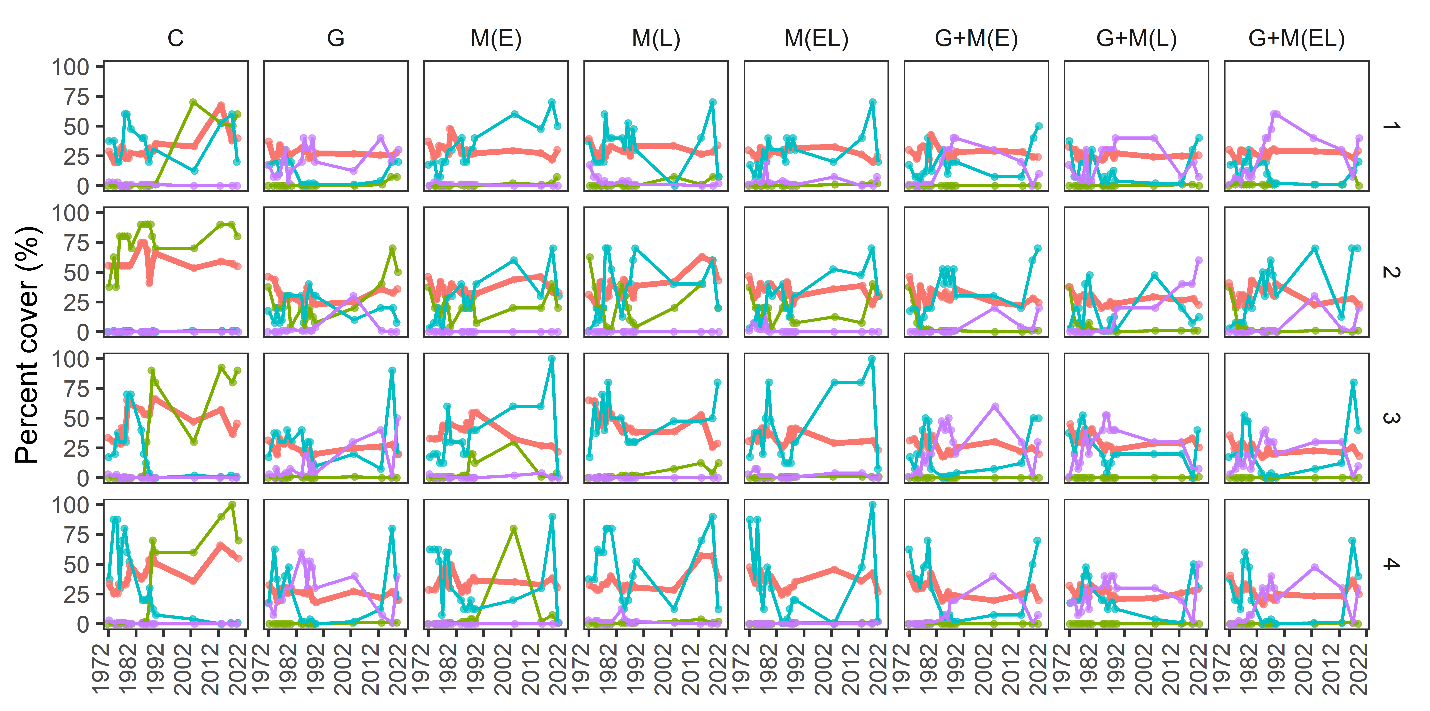

**Fig. S2.** **Dominance (thick red lines), percent cover of *Elytrigia atherica* (yellowgreen), *Festuca rubra* (light blue), and *Juncus gerardii* (purple) in four blocks of different treatments over the 48-year experiment.** Four blocks were established in 1972, encompassing different plant communities characterized by different dominant species: block 1) *F. rubra* and *Armeria maritima*; block 2) *E. atherica*; block 3) *F. rubra* and *Artemisia maritima*; block 4) *F. rubra* and *Limonium vulgare*. Block 1 and 2 were situated in high marsh, block 3, and 4 in the low marsh. The treatment description corresponds to that of Fig. S1.

Table S1. List of species occurring during the 48-year experiment and their life form. Species in boldface belong to the halophytes, according to Scherfose (1990) and Bakker *et al.* (2002). Life form was classified according to https://wilde-planten.nl.

| Species name | Life form |
| --- | --- |
| *Agrostis stolonifera* | graminoid |
| *Armeria maritima* | forb |
| ***Artemisia maritima*** | woody |
| ***Aster tripolium*** | forb |
| ***Atriplex littoralis*** | forb |
| ***Atriplex portulacoides*** | woody |
| ***Atriplex prostrata*** | forb |
| *Bromus hordeaceus ssp. hordeaceus* | graminoid |
| *Bupleurum tenuissimum* | forb |
| *Carex distans* | graminoid |
| *Carex extensa* | graminoid |
| *Centaurium littorale* | forb |
| *Centaurium pulchellum* | forb |
| *Cerastium fontanum ssp. vulgare* | forb |
| *Cirsium arvense* | forb |
| *Cochlearia danica* | forb |
| ***Cochlearia officinalis ssp. anglica*** | forb |
| *Elytrigia atherica* | graminoid |
| *Elytrigia repens* | graminoid |
| *Festuca rubra* | graminoid |
| ***Glaux maritima*** | forb |
| *Hippophae rhamnoides* | woody |
| *Holcus lanatus* | graminoid |
| *Honckenya peploides* | forb |
| ***Juncus gerardii*** | graminoid |
| *Juncus maritimus* | graminoid |
| *Leontodon autumnalis* | forb |
| ***Limonium vulgare*** | forb |
| *Lolium perenne* | graminoid |
| *Lotus corniculatus* | legume |
| *Odontites vernus* ssp*. serotinus* | forb |
| *Parapholis strigosa* | graminoid |
| *Plantago coronopus* | forb |
| *Plantago lanceolata* | forb |
| *Plantago major* | forb |
| ***Plantago maritima*** | forb |
| *Poa annua* | graminoid |
| *Poa pratensis* | graminoid |
| *Potentilla anserina* | forb |
| *Prunus serotina* | woody |
| ***Puccinellia maritima*** | graminoid |
| *Puccinellia rupestris* | graminoid |
| *Rumex crispus* | forb |
| ***Sagina maritima*** | forb |
| *Sagina nodosa* | forb |
| *Sagina procumbens* | forb |
| ***Salicornia* spp.** | forb |
| ***Spergularia* spp*.*** | forb |
| *Stellaria graminea* | forb |
| *Stellaria media* | forb |
| ***Suaeda maritima*** | forb |
| *Taraxacum* spp. | forb |
| *Trifolium fragiferum* | forb |
| *Trifolium pratense* | legume |
| *Trifolium repens* | legume |
| ***Triglochin maritima*** | graminoid |

Table S2. Temporal trends (smooths) estimated from the generalized additive mixed models for plant diversity at the local and landscape scales.

| Variable | Smooths | edf | F | p-value | R^2^ | s.d. | phi |
| --- | --- | --- | --- | --- | --- | --- | --- |
| Plant diversity in local communities | s(year):treatmentC | 1.13 | 5.22 | 0.0159 | 0.50 | 0.22 | 0.58 |
|  | s(year):treatmentG | 2.75 | 3.1 | 0.0849 |  |  |  |
|  | s(year):treatmentM(E) | 1 | 0.12 | 0.7284 |  |  |  |
|  | s(year):treatmentM(L) | 1.17 | 4.87 | 0.0312 |  |  |  |
|  | s(year):treatmentM(EL) | 2.32 | 2.31 | 0.1652 |  |  |  |
|  | s(year):treatmentG+M(E) | 4 | 5 | 0.0007 |  |  |  |
|  | s(year):treatmentG+M(L) | 3.06 | 2.43 | 0.0566 |  |  |  |
|  | s(year):treatmentG+M(EL) | 2.92 | 2.29 | 0.1366 |  |  |  |
| Plant diversity in the landscape | s(year):treatmentC | 1 | 20.12 | < 0.0001 | 0.71 | NA | 0.40 |
|  | s(year):treatmentG | 1 | 1.06 | 0.3058 |  |  |  |
|  | s(year):treatmentM(E) | 1 | 0.21 | 0.6474 |  |  |  |
|  | s(year):treatmentM(L) | 1.72 | 2.78 | 0.0438 |  |  |  |
|  | s(year):treatmentM(EL) | 1.65 | 0.39 | 0.5689 |  |  |  |
|  | s(year):treatmentG+M(E) | 3.53 | 2.61 | 0.0297 |  |  |  |
|  | s(year):treatmentG+M(L) | 1 | 0.24 | 0.6257 |  |  |  |
|  | s(year):treatmentG+M(EL) | 1 | 0.37 | 0.5439 |  |  |  |

edf: effective degrees of freedom, the larger the values, the wiggiler the trends. s.d.: the standard errors for random variable (block). Phi indicates how quickly the correlation between any two residuals decreases as a function of their time difference.

Table S3. Temporal trends (smooths) estimated from the generalized additive mixed models for plant diversity in different functional groups at the local and landscape scales.

| Variable | Smooths | edf | Ref.df | F | p-value | R^2^ | s.d. | phi |
| --- | --- | --- | --- | --- | --- | --- | --- | --- |
| Number of forbs in local communities | s(year):treatmentC | 1.89 | 1.89 | 1 | 0.28 | 0.48 | 0.30 | 0.38 |
|  | s(year):treatmentG | 2.3 | 2.3 | 2.58 | 0.1234 |  |  |  |
|  | s(year):treatmentM(E) | 1 | 1 | 4.51 | 0.0342 |  |  |  |
|  | s(year):treatmentM(L) | 1.46 | 1.46 | 0.23 | 0.7714 |  |  |  |
|  | s(year):treatmentM(EL) | 1.14 | 1.14 | 3.01 | 0.0644 |  |  |  |
|  | s(year):treatmentG+M(E) | 3.72 | 3.72 | 4.88 | 0.0008 |  |  |  |
|  | s(year):treatmentG+M(L) | 3.67 | 3.67 | 3.93 | 0.0043 |  |  |  |
|  | s(year):treatmentG+M(EL) | 2.87 | 2.87 | 2.45 | 0.1402 |  |  |  |
| Number of graminoids in local communities | s(year):treatmentC | 1.65 | 1.65 | 4.11 | 0.0139 | 0.37 | 0.38 | 0.66 |
|  | s(year):treatmentG | 2.74 | 2.74 | 3.32 | 0.066 |  |  |  |
|  | s(year):treatmentM(E) | 1.25 | 1.25 | 2.85 | 0.1123 |  |  |  |
|  | s(year):treatmentM(L) | 1 | 1 | 8.53 | 0.0036 |  |  |  |
|  | s(year):treatmentM(EL) | 2.95 | 2.95 | 2.86 | 0.0622 |  |  |  |
|  | s(year):treatmentG+M(E) | 3.14 | 3.14 | 2.99 | 0.024 |  |  |  |
|  | s(year):treatmentG+M(L) | 1 | 1 | 0.05 | 0.8167 |  |  |  |
|  | s(year):treatmentG+M(EL) | 2.3 | 2.3 | 2.2 | 0.1573 |  |  |  |
| Number of halophytes in local communities | s(year):treatmentC | 1 | 1 | 3.57 | 0.0593 | 0.35 | 1.90 | 0.60 |
|  | s(year):treatmentG | 2.93 | 2.93 | 4.68 | 0.0074 |  |  |  |
|  | s(year):treatmentM(E) | 1 | 1 | 1.1 | 0.2945 |  |  |  |
|  | s(year):treatmentM(L) | 1.87 | 1.87 | 1.18 | 0.3741 |  |  |  |
|  | s(year):treatmentM(EL) | 1.58 | 1.58 | 0.29 | 0.6298 |  |  |  |
|  | s(year):treatmentG+M(E) | 3.24 | 3.24 | 4.51 | 0.0029 |  |  |  |
|  | s(year):treatmentG+M(L) | 3.34 | 3.34 | 5.02 | 0.002 |  |  |  |
|  | s(year):treatmentG+M(EL) | 2.45 | 2.45 | 2.31 | 0.1999 |  |  |  |
| Number of forbs in the landscape | s(year):treatmentC | 1 | 1 | 1.44 | 0.2326 | 0.67 | NA | 0.39 |
|  | s(year):treatmentG | 1 | 1 | 7.23 | 0.0081 |  |  |  |
|  | s(year):treatmentM(E) | 1 | 1 | 3.06 | 0.0827 |  |  |  |
|  | s(year):treatmentM(L) | 2.88 | 2.88 | 1.92 | 0.1855 |  |  |  |
|  | s(year):treatmentM(EL) | 1 | 1 | 4.75 | 0.0312 |  |  |  |
|  | s(year):treatmentG+M(E) | 3.65 | 3.65 | 2.03 | 0.0616 |  |  |  |
|  | s(year):treatmentG+M(L) | 1 | 1 | 0.42 | 0.5203 |  |  |  |
|  | s(year):treatmentG+M(EL) | 1.44 | 1.44 | 0.79 | 0.5511 |  |  |  |
| Number of graminoids in the landscape | s(year):treatmentC | 2.79 | 2.79 | 36.55 | 0.0000 | 0.77 | NA | NA |
|  | s(year):treatmentG | 3.97 | 3.97 | 3.98 | 0.0035 |  |  |  |
|  | s(year):treatmentM(E) | 1 | 1 | 13.38 | 0.0004 |  |  |  |
|  | s(year):treatmentM(L) | 1 | 1 | 58.11 | 0.0000 |  |  |  |
|  | s(year):treatmentM(EL) | 6.3 | 6.3 | 6.61 | 0.0000 |  |  |  |
|  | s(year):treatmentG+M(E) | 3.02 | 3.02 | 5.25 | 0.0019 |  |  |  |
|  | s(year):treatmentG+M(L) | 1 | 1 | 5.65 | 0.0191 |  |  |  |
|  | s(year):treatmentG+M(EL) | 1 | 1 | 0.34 | 0.5610 |  |  |  |
| Number of halophytes in the landscape | s(year):treatmentC | 1 | 1 | 8.38 | 0.0045 | 0.75 | NA | NA |
|  | s(year):treatmentG | 2.15 | 2.15 | 2.83 | 0.0747 |  |  |  |
|  | s(year):treatmentM(E) | 1 | 1 | 2.54 | 0.1134 |  |  |  |
|  | s(year):treatmentM(L) | 5.22 | 5.22 | 2.61 | 0.0196 |  |  |  |
|  | s(year):treatmentM(EL) | 2.33 | 2.33 | 2.9 | 0.0348 |  |  |  |
|  | s(year):treatmentG+M(E) | 4 | 4 | 5.61 | 0.0004 |  |  |  |
|  | s(year):treatmentG+M(L) | 2.26 | 2.26 | 2.27 | 0.1726 |  |  |  |
|  | s(year):treatmentG+M(EL) | 2.51 | 2.51 | 4.37 | 0.0183 |  |  |  |

edf: effective degrees of freedom, the larger the values, the wiggiler the trends. s.d.: the standard errors for random variable (block). Phi indicates how quickly the correlation between any two residuals decreases as a function of their time difference. Autocorrelation was not included in the models for number of graminoids in the landscape and number of halophytes in the landscape, because in both cases ΔAIC < 4 for models with and without autocorrelation structure.

Table S4. Temporal trends (smooths) estimated from the generalized additive mixed models for temporal and spatial community dissimilarity and their components.

| Variable | Smooths | edf | Ref.df | F | p-value | R^2^ | s.d. | phi |
| --- | --- | --- | --- | --- | --- | --- | --- | --- |
| Temporal community dissimilarity | s(year):treatmentC | 1 | 1 | 0.01 | 0.9063 | 0.18 | 0.00 | 0.31 |
|  | s(year):treatmentG | 1 | 1 | 10.46 | 0.0013 |  |  |  |
|  | s(year):treatmentM(E) | 2.4 | 2.4 | 7.57 | 0.0002 |  |  |  |
|  | s(year):treatmentM(L) | 1 | 1 | 8.72 | 0.0033 |  |  |  |
|  | s(year):treatmentM(EL) | 2.58 | 2.58 | 4.99 | 0.0026 |  |  |  |
|  | s(year):treatmentG+M(E) | 1.83 | 1.83 | 1.51 | 0.3131 |  |  |  |
|  | s(year):treatmentG+M(L) | 2.29 | 2.29 | 4.37 | 0.0085 |  |  |  |
|  | s(year):treatmentG+M(EL) | 2.58 | 2.58 | 6.08 | 0.0009 |  |  |  |
| Temporal balanced variation | s(year):treatmentC | 2.24 | 2.24 | 2.53 | 0.057 | 0.24 | 0.01 | 0.18 |
|  | s(year):treatmentG | 1 | 1 | 21.14 | < 0.0001 |  |  |  |
|  | s(year):treatmentM(E) | 4.85 | 4.85 | 4.38 | 0.0005 |  |  |  |
|  | s(year):treatmentM(L) | 1.9 | 1.9 | 2.66 | 0.0467 |  |  |  |
|  | s(year):treatmentM(EL) | 3.62 | 3.62 | 6.84 | 0.0001 |  |  |  |
|  | s(year):treatmentG+M(E) | 2.23 | 2.23 | 6.11 | 0.0029 |  |  |  |
|  | s(year):treatmentG+M(L) | 2.15 | 2.15 | 4.78 | 0.0075 |  |  |  |
|  | s(year):treatmentG+M(EL) | 2.93 | 2.93 | 11.93 | < 0.0001 |  |  |  |
| Temporal abundance gradient | s(year):treatmentC | 2.43 | 2.43 | 6.03 | 0.0028 | 0.20 | 0.03 | NA |
|  | s(year):treatmentG | 1 | 1 | 2.27 | 0.1324 |  |  |  |
|  | s(year):treatmentM(E) | 5.27 | 5.27 | 4.6 | 0.0003 |  |  |  |
|  | s(year):treatmentM(L) | 2.25 | 2.25 | 3.94 | 0.0177 |  |  |  |
|  | s(year):treatmentM(EL) | 3.39 | 3.39 | 2.19 | 0.0938 |  |  |  |
|  | s(year):treatmentG+M(E) | 5.38 | 5.38 | 5.57 | 0 |  |  |  |
|  | s(year):treatmentG+M(L) | 1.32 | 1.32 | 0.15 | 0.6463 |  |  |  |
|  | s(year):treatmentG+M(EL) | 6.09 | 6.09 | 3.68 | 0.0009 |  |  |  |
| Spatial community dissimilarity | s(year):treatmentC | 1 | 1 | 20.71 | < 0.0001 | 0.38 | NA | 0.65 |
|  | s(year):treatmentG | 1 | 1 | 0.08 | 0.7818 |  |  |  |
|  | s(year):treatmentM(E) | 2.76 | 2.76 | 2.87 | 0.0448 |  |  |  |
|  | s(year):treatmentM(L) | 2.21 | 2.21 | 1.23 | 0.3606 |  |  |  |
|  | s(year):treatmentM(EL) | 4.8 | 4.8 | 1.53 | 0.1253 |  |  |  |
|  | s(year):treatmentG+M(E) | 1 | 1 | 3.66 | 0.0582 |  |  |  |
|  | s(year):treatmentG+M(L) | 1.79 | 1.79 | 0.69 | 0.4047 |  |  |  |
|  | s(year):treatmentG+M(EL) | 1 | 1 | 0.46 | 0.4968 |  |  |  |
| Spatial balanced variation | s(year):treatmentC | 1 | 1 | 14 | 0.0003 | 0.24 | NA | 0.63 |
|  | s(year):treatmentG | 1 | 1 | 0.64 | 0.4269 |  |  |  |
|  | s(year):treatmentM(E) | 2.57 | 2.57 | 2.55 | 0.0706 |  |  |  |
|  | s(year):treatmentM(L) | 1.84 | 1.84 | 0.68 | 0.4659 |  |  |  |
|  | s(year):treatmentM(EL) | 1.66 | 1.66 | 0.29 | 0.6188 |  |  |  |
|  | s(year):treatmentG+M(E) | 1 | 1 | 2.14 | 0.1464 |  |  |  |
|  | s(year):treatmentG+M(L) | 1.3 | 1.3 | 0.11 | 0.8305 |  |  |  |
|  | s(year):treatmentG+M(EL) | 1 | 1 | 0.06 | 0.8044 |  |  |  |
| Spatial abundance gradient | s(year):treatmentC | 1 | 1 | 0.17 | 0.6847 | 0.12 | NA | 0.28 |
|  | s(year):treatmentG | 1 | 1 | 4.16 | 0.0435 |  |  |  |
|  | s(year):treatmentM(E) | 1 | 1 | 0.12 | 0.733 |  |  |  |
|  | s(year):treatmentM(L) | 1 | 1 | 0.48 | 0.4917 |  |  |  |
|  | s(year):treatmentM(EL) | 1 | 1 | 0.29 | 0.5897 |  |  |  |
|  | s(year):treatmentG+M(E) | 1 | 1 | 1.04 | 0.3103 |  |  |  |
|  | s(year):treatmentG+M(L) | 1 | 1 | 2.39 | 0.1242 |  |  |  |
|  | s(year):treatmentG+M(EL) | 1 | 1 | 1.7 | 0.1952 |  |  |  |

edf: effective degrees of freedom, the larger the values, the wiggiler the trends. s.d.: the standard errors for random variable (block). Phi indicates how quickly the correlation between any two residuals decreases as a function of their time difference. Autocorrelation was not included in the model of temporal abundance gradient, because ΔAIC < 4 for models with and without autocorrelation structure.

**References**

Bakker, J.P., Esselink, P., Dijkema, K.S., Van Duin, W.E. & De Jong, D.J. (2002). Restoration of salt marshes in the Netherlands. *Hydrobiologia*, 478, 29–51.

Guevara, M. R., Hartmann, D., & Mendoza, M. (2016). diverse: an R Package to Measure

Diversity in Complex Systems. The R Journal, 8(2), 60–78. URL https://journal.r-project.org/archive/2016-2/guevara-hartmann-mendoza.pdf.

Koerner, S.E., Smith, M.D., Burkepile, D.E., Hanan, N.P., Avolio, M.L., Collins, S.L., *et al.* (2018). Change in dominance determines herbivore effects on plant biodiversity. *Nat. Ecol. Evol.*, 2, 1925–1932.

Kuznetsova A, Brockhoff PB, Christensen RHB (2017). “lmerTest Package: Tests in Linear Mixed Effects Models.” *Journal of Statistical Software*, **82**(13), 1–26. doi: [10.18637/jss.v082.i13](https://doi.org/10.18637/jss.v082.i13).

Scherfose, V. (1990) Salz‐Zeigerwerte von Gefäßpflanzen der Salzmarschen, Tideröhrichte und Salzwassertümpel an der deutschen Nord‐ und Ostseeküste. Jahrbuch Niedersächsisches Landesamt Wasser und Abfall, Forschungsstelle Küste, 39, 31–82.
